# Pacific Biosciences long reads-based genome sequencing data from a widespread bee fungal parasite, *Nosema ceranae*

**DOI:** 10.1101/2020.04.05.026849

**Authors:** Huazhi Chen, Wende Zhang, Yu Du, Xiaoxue Fan, Jie Wang, Haibin Jiang, Yuanchan Fan, Zhiwei Zhu, Cuiling Xiong, Yanzhen Zheng, Dafu Chen, Rui Guo

**Affiliations:** College of Animal Sciences (College of Bee Science), Fujian Agriculture and Forestry University, Fuzhou 350002, China

**Keywords:** Third-generation sequencing, PacBio, Long reads, Honeybee, *Nosema ceranae*, genome

## Abstract

*Nosema ceranae* is a widespread fungal parasite that infects both adult honeybee and honeybee larvae, leading to microsporidiosis, which seriously affects bee health and apicultural industry. In this article, genome sequencing of clean spores of *N. ceranae* was conducted using third-generation Pacific Biosciences (PacBio) single molecule real time (SMRT) sequencing technology. In total, 152671 subreads were obtained after quality control of raw reads from PacBio SMRT sequencing, with a N50 and average length of 14422 bp and 11310 bp, respectively. Additionally, the length distribution of subreads was from 10000 bp to more than 50000 bp. Nineteen scaffords with a total length of 7354221 bp were assembled, and the N50, N90 and maximum scafford length were 728543 bp, 198795 bp and 1917792 bp, respectively. The GC content was 25.97%. Furthermore, by integration of genes predicted from *de novo* and homology-based methods, 3112 *N. ceranae* genes were finally assembled, with a total length of 2730179 bp and mean length of 877.31 bp. In addition, the total length and mean length of exons were 2657637 bp and 854 bp, respectively; and the total length and mean length of introns were 72542 bp and 23.31 bp, respectively. The genome sequencing data documented here will give deep insights into the molecular biology of *N. ceranae*, facilitate exploration of genes and pathways associated with toxin factors and infection-related factors, and benefit research on comparative genomics and phylogenetic diversity of *Nosema* species.

## Value of the data

This genome dataset gives deep insights into the molecular biology of *N. ceranae*, which is beneficial for researchers working on bee microsporidiosis.

The data could be used to study *N. ceranae* genes and pathways relevant to toxin factors and infection-related factors.

Our genome data is valuable for the scientific community to perform study on comparative genomics and investigate phylogenetic diversity of *Nosema* species.

Current genome data sheds light on the development of novel control strategies for bee microsporidiosis caused by *N. ceranae*.

## Data

The data reported here was derived from genome sequencing of *N. ceranae* spores using Pacific Biosciences (PacBio) single molecule real time (SMRT) sequencing technology. A total of 152671 subreads were obtained after quality control of raw reads generated from *N. ceranae* spores, with a N50 and average length of 14422 bp and 11310 bp, respectively (**Table 1**). Additionally, the length distribution of subreads was from 10000 bp to more than 50000 bp (**Figure 1**). Using a combination of Canu and wtbdg softwares, 19 scaffords were assembled, with a total length of 7354221 bp (**Table 2**). The N50, N90 and maximum scafford length were 728543 bp, 198795 bp and 1917792 bp, respectively (**Table 2**). Meanwhile, there were 36 contigs with a total length of 7318568 bp, and the N50, N90 and maximum contig length were 457301 bp, 96012 bp and 824658 bp, respectively (**Table 2**). Besides, the GC content was 25.97% (**Table 2**). Two different methods were used to conduct gene prediction, including *de novo* prediction and homology-based prediction. Based on former approach, 2361, 3025 and 3223 genes were respectively predicted using Augustus, GlimmerHMM and SNAP (**Table 3**). On basis of homology-based prediction method, 1615 genes homologous to *Nosem apis* BRLO1, 1653 genes homologous to *Nosema bombycis* CQ1, and 3969 genes homologous to *N. ceranae* PA08 1199 were predicted (**Table 3**). Further, 3112 *N. ceranae* genes were assembled by integration of results from above-mentioned two methods (**Table 3**). As shown in **Table 4**, the total length of *N. ceranae* genes was 2730179 bp, and the mean length was 877.31 bp. As shown in **Table 4**, the total length and average length of exons were 2657637 bp and 854 bp, respectively; and the total length and average length of introns were 72542 bp and 23.31 bp, respectively.

**Table 1.**
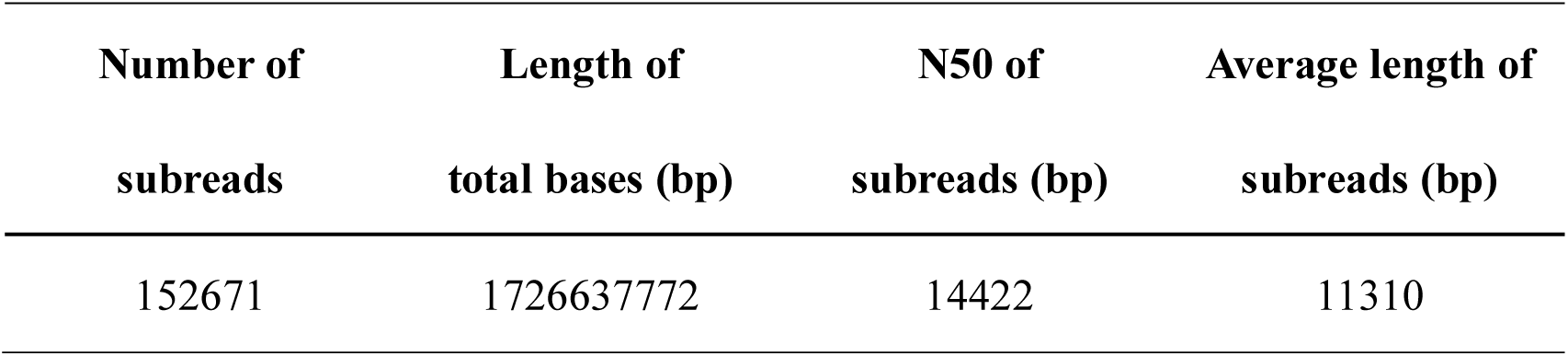
Overview of subreads after quality control of raw reads generated from PacBio SMRT sequencing of *N. ceranae* spore.

| Number of<br>subreads | Length of<br>total bases (bp) | N50 of<br>subreads (bp) | Average length of<br>subreads (bp) |
| --- | --- | --- | --- |
| 152671 | 1726637772 | 14422 | 11310 |

**Table 2.** Assembly statistics of *N. ceranae* genome.

| Attributes | Value |
| --- | --- |
| Scaffold number | 19 |
| Scaffold length (bp) | 7354221 |
| Scaffold N50 (bp) | 728543 |
| Scaffold N90 (bp) | 198795 |
| Maximum scaffold (bp) | 1917792 |
| Total length of gaps (bp) | 35653 |
| Contig number | 36 |
| Contig length (bp) | 7318568 |
| Contig N50 (bp) | 457301 |
| Contig N90 (bp) | 96012 |
| Maximum contig (bp) | 824658 |
| GC content (%) | 25.97 |

**Table 3.**
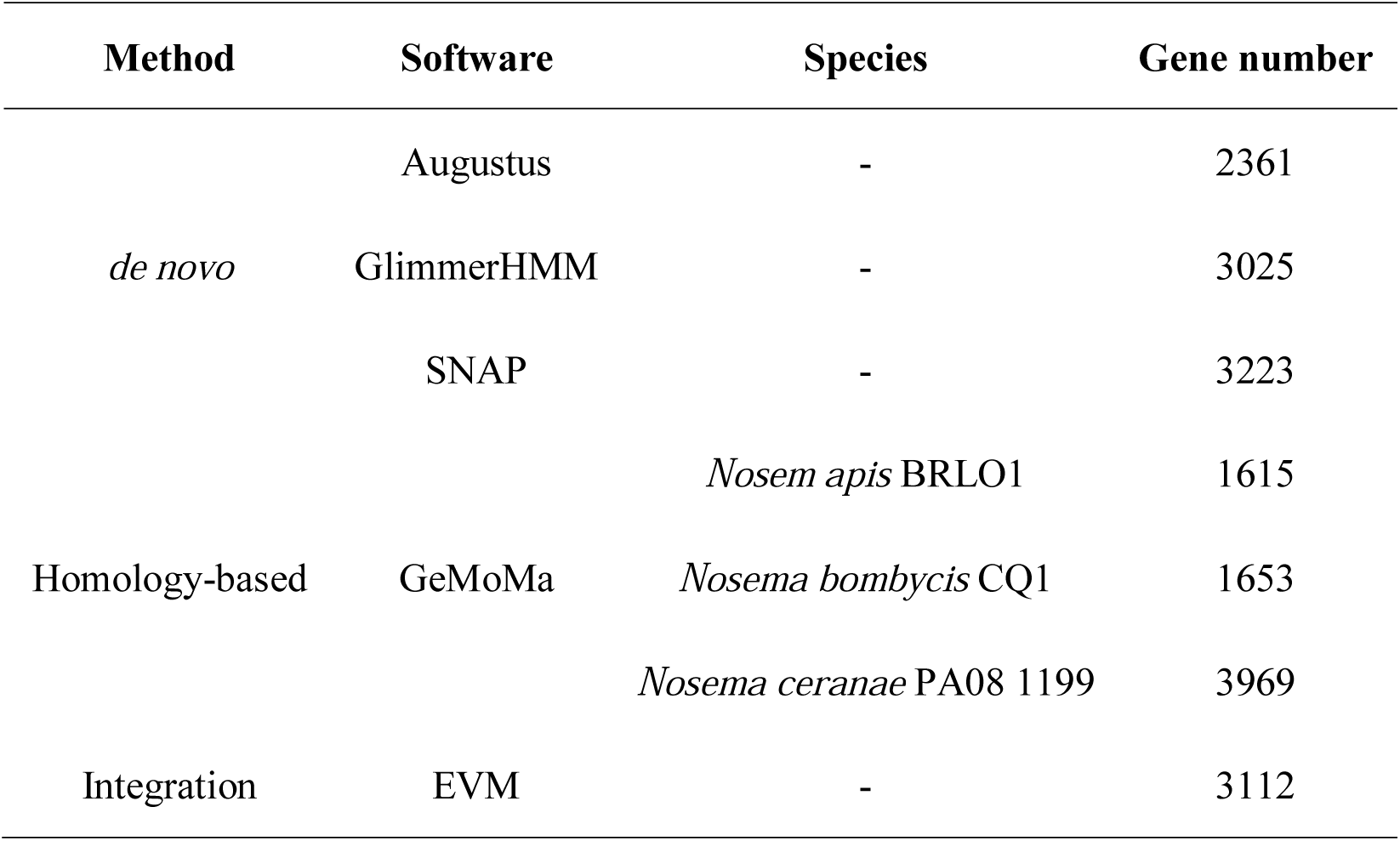
Summary of gene prediction based on various bioinformatics softwares.

**Table 4.** Genome features of *Nosema ceranae*.

| Gene number | Gene length (bp) | Average gene length (bp) | Exon length (bp) | Average exon length (bp) | Intron length (bp) | Average intron length (bp) |
| --- | --- | --- | --- | --- | --- | --- |
| 3112 | 2730179 | 877.31 | 2657637 | 854 | 72542 | 23.31 |

**Figure 1.**
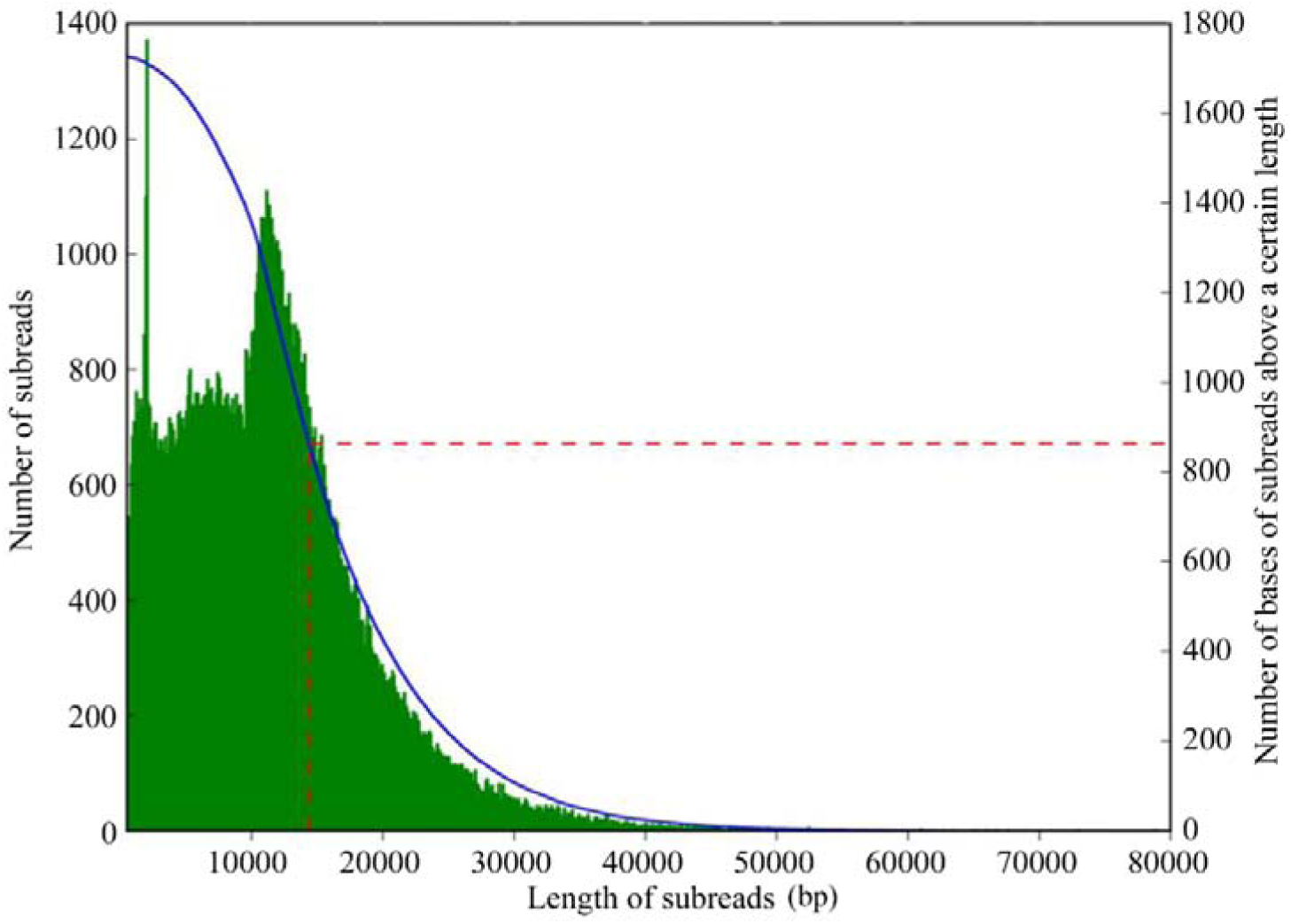
Length distribution of subreads.

## Experimental Design, Materials, and Methods

### Fungal spore preparation

Following the method described by Cornman et al. [1] with some minor modifications [2], using Percoll discontinuous density gradient centrifugation, *N. ceranae* spores were previously purified from an *N. ceranae*-infected western honeybee colony at College of Animal Sciences (College of Bee Science) in Fujian Agriculture and Forestry University. A bit of spores was examined via PCR and confirmed to be mono-specific [3] with previously developed specific primers [4]. The clean spores of *N. ceranae* were immediately frozen in liquid nitrogen and stored at - 80 °C until genome sequencing.

### Illumina short reads sequencing

The 270 bp library was constructed according to Illumina’s standard protocol, including fragmentation of genomic DNA, end repair, adaptor ligation and PCR amplification. Subsequently, the 270 bp library was quantified using 2100 Bioanalyzer (Agilent, USA) and subjected to paired-ended 150 bp sequencing on an Illumina HiSeq X Ten platform. The second-generation sequencing data (filtered reads: 1.30 Gb, sequencing depth: 177.75×) were used to estimate the genome size, repeat content, and heterozygosity.

### PacBio SMRT long reads sequencing and genome assembly

Firstly, the 10 kb library was constructed following PacBio’s standard instruction, including fragmentation of genomic DNA, end repair, adaptor ligation, and templates purification. Secondly, the 10-kb library was quantified by 2100 Bioanalyzer (Agilent, USA) and sequenced using PacBio SMRT [5] by Biomarker Technologies (Beijing, China), and the third-generation sequencing data (filtered reads: 1.73 Gb, sequencing depth: 234.78×) was assembled by CANU (Version-1.2) with default parameters [6]. Finally, Illumina reads were used for error correction and gap filling with SOAPdenovo GAPCLOSER v1.12 [7].

## Acknowledgments

This research was financially supported by the Earmarked Fund for China Agriculture Research System (No. CARS-44-KXJ7), the Science and Technology Planning Project of Fujian Province (No. 2018J05042), the Teaching and Scientific Research Fund of Education Department of Fujian Province (No. JAT170158), the Outstanding Scientific Research Manpower Fund of Fujian Agriculture and Forestry University (No. xjq201814), and the Scientific and Technical Innovation Fund of Fujian Agriculture and Forestry University (No. CXZX2017342, No. CXZX2017343).

